# Improving Protein Docking with Constraint Programming and Coevolution Data

**DOI:** 10.1101/002329

**Authors:** Ludwig Krippahl, Fábio Madeira

## Abstract

**Background:** Constraint programming (CP) is usually seen as a rigid approach, focusing on crisp, precise, distinctions between what is allowed as a solution and what is not. At first sight, this makes it seem inadequate for bioinformatics applications that rely mostly on statistical parameters and optimization. The prediction of protein interactions, or protein docking, is one such application. And this apparent problem with CP is particularly evident when constraints are provided by noisy data, as it is the case when using the statistical analysis of Multiple Sequence Alignments (MSA) to extract coevolution information. The goal of this paper is to show that this first impression is misleading and that CP is a useful technique for improving protein docking even with data as vague and noisy as the coevolution indicators that can be inferred from MSA.

**Results:** Here we focus on the study of two protein complexes. In one case we used a simplified estimator of interaction propensity to infer a set of five candidate residues for the interface and used that set to constrain the docking models. Even with this simplified approach and considering only the interface of one of the partners, there is a visible focusing of the models around the correct configuration. Considering a set of 400 models with the best geometric contacts, this constraint increases the number of models close to the target (RMSD ¡5Å) from 2 to 5 and decreases the RMSD of all retained models from 26Å to 17.5Å. For the other example we used a more standard estimate of coevolving residues, from the Co-Evolution Analysis using Protein Sequences (CAPS) software. Using a group of three residues identified from the sequence alignment as potentially co-evolving to constrain the search, the number of complexes similar to the target among the 50 highest scoring docking models increased from 3 in the unconstrained docking to 30 in the constrained docking.

**Conclusions:** Although only a proof-of-concept application, our results show that, with suitably designed constraints, CP allows us to integrate coevolution data, which can be inferred from databases of protein sequences, even though the data is noisy and often “fuzzy”, with no well-defined discontinuities. This also shows, more generally, that CP in bioinformatics needs not be limited to the more crisp cases of finite domains and explicit rules but can also be applied to a broader range of problems that depend on statistical measurements and continuous data.

## 1. INTRODUCTION

Proteins constitute the main machinery of life. These macromolecules, consisting of long strings of amino acid residues, with tens or hundreds of thousands of atoms, interact in complex ways to regulate and catalyse the chemistry of living organisms. In some cases, interactions occur because, by chance, the sequence and structure of a protein makes it recognize a specific substrate. This happens in the immune system, for example, where antibodies are created by random shuffling of genetic motifs. But, in general, proteins interact because they evolved together under pressure to optimize a particular interaction, and this coevolution can leave traces in the sequences of the proteins involved (Pazos et al., 1997). For two proteins to interact in a biologically useful manner they must recognize each other and form a complex that is stable enough to promote a particular effect, be it a chemical reaction, electron or substrate transfer, or conformational change. This requires that the interface region of one protein matches, in some chemical or physical attributes, the interface region of the other protein, thus placing evolutionary constraints on the residues involved. A mutation in one of these interface residues will create a selection pressure favouring compensatory mutations in the other protein (Pazos and Valencia, 2008).

This is also true for those interactions, within each protein, that determine its three-dimensional structure. Thus, identifying coevolution of protein residues is of great potential interest for structure prediction (Codoñer and Fares, 2008), protein domain analysis (Yeang and Haussler, 2007), and for the study of the relations between structure and function (Little and Chen, 2009). However, coevolution data is insufficient for determining the structure of a protein. The statistical information on correlations or mutual information between residues is too noisy to provide more than a few clear signals, and knowing only a few contacts is not enough to determine the structure of a protein. Furthermore, correlation between residues can arise from other factors besides the pressure to maintain the necessary contacts. Residues can be correlated because unrelated ancestral mutations persist in the same descendant lineages, or because of functional or structural constraints that affect both residues even if they are not close together in the structure (Lovell and Robertson, 2010). In addition, paralogous sequences – those that descend from a common ancestor in the same organism lineage by gene duplication – can diverge by selection for different roles (e.g. myoglobin and hemoglobin). Since it is not trivial to distinguish paralogs from orthologs – those that retain the same role and diverge due to the splitting of the organism lineages – this interferes with the statistics obtained from the MSA.

For protein interaction prediction, the scarcity of contact information should, in theory, not be such a problem in itself. Since the structure of each interaction partner is known, identifying even a single correct contact would greatly restrict the number of possible configurations for the complex. However, the problem remains that any correct contact is likely to be hidden in a set of incorrect candidates, because correlations can arise from stochastic factors or from evolutionary factors unrelated to interface contacts. For these reasons, there are many different ways of trying to calculate the appropriate correlation, and the definition of what measure is the best for each case is still an open problem (see (Halperin et al., 2006) and (Horner et al., 2007) for an overview of several alternatives), but our focus, in this paper, is not to address the detailed issues of how to infer inter-molecular contacts from MSA data. In fact, our measure is a simplification of different statistical correlation measures and scores based on evolutionary information and contact propensities (e.g. (Madaoui and Guerois, 2008)). Rather, our focus is to show that despite the nature of the problem, and of the data behind the constraints, CP can be used to integrate this information and improve protein docking predictions. We achieve this by using a constrained docking algorithm that is robust to noisy data.

### 1.1 Constrained docking

As published in previous work, BiGGER (Palma et al, 2000), our protein docking application, can model a wide range of geometric constraints on the contacts between two proteins by imposing a cardinality constraint on the set of contacts between two specified groups of points. Given two groups, *A* and *B*, with *M*_*a*_ and *M*_*b*_ points respectively, and with each group belonging to each partner, the constraint is that there must be at least *N* contacts between the two groups. A contact is defined as a pair of points (*P*_*a*_; *P*_*b*_), respectively from groups *A* and *B*, that have a distance below a predefined contact threshold *D*. As long as *N* is not larger than the number of correct contacts between these groups and *D* is not too restrictive, the search space can be reduced while still avoiding inconsistencies due to having incorrect residues included in the groups or losing correct models due to over-constraining the search. The details for this constraint, and how it is propagated during docking, can be found in (Krippahl and Barahona, 2005). Moreover, we showed that even a few contacts are enough to pinpoint the correct protein complex, assuming that such information is available (Krippahl et al., 2003), and that this CP-based can improve docking the results when used in conjunction with specific information about the mechanism of interaction (Monaco et al., 2007) or with spectroscopic data (Palma et al., 2005). However, it is still an open question whether one can, in practice, apply this same technique to less accurate data. Hence, the importance of trying to combine our constrained docking algorithm with statistical indicators on coevolution.

### 1.2 Multiple Sequence Alignment

Another problem with the inference of coevolutionary data is the MSA itself. To identify coevolving residues it is first necessary to compute an MSA over a set of similar sequences, in order to compare the mutations at different positions. The MSA attempts to represent the evolutionary relations between different sequences by aligning them under the assumption that mutations are independent, and is usually depicted as a table where each row represents one sequence and each column consists of all residues that descend from a common ancestor at that position, in some ancestral sequence. Deletions or insertions are represented by adding gaps to the appropriate sequences. Since computing a MSA is an NP-Hard problem ((Wang and Jiang, 1994), (Just, 2001), (Elias, 2006)), MSA algorithms need to take shortcuts and rely heavily on heuristics trying to optimize the alignment score. This often leads to poor alignments and, given the importance of this problem, there has been a significant effort to find ways of improving MSA by modelling constraints within the classical alignment algorithms ((Myers et al., 1996), (Sammeth et al., 2003), (Papadopoulos and Agarwala, 2007)) or with Constraint Programming techniques ((Yap, 2001), (Will et al., 2008)). However, one important problem for identifying coevolution is the alignment score itself. Alignments are scored by assuming independent mutations and estimating the likelihood of each pair of matching residues, but if some positions are linked by coevolutionary constraints, scoring their alignments independently will not reflect their actual evolutionary relations. While attempting to align residues that minimize the differences between sequences, the MSA algorithm can misplace coevolving sites in the alignment, thus making it harder to find the statistical correlations that indicate coevolution. However, the possibility of specifying groups of atoms that are in contact without having to specify which particular pairs form such contacts, a feature of our cardinality constraint over the set of contacts, allows us to improve the reliability of the information inferred from the sequence alignments by averaging the estimates for each residue, as detailed below in the methods section.

## 2. METHODS

Our method spans the complete docking process, starting from the structures of the proteins that interact and finishing with a prediction for the complex. The first step is to use the sequence of each interaction partner to query a database for similar sequences. In our case, we used the EBI PSI-BLAST server. The reason for using PSI-BLAST (Altschul et al., 1997) instead of a simple BLAST search is that we need to start with a large number of homologous sequences, and PSI-BLAST allows us to find more distant relatives. Once we obtain a large set of homologous sequences, we use ClustalW (Chenna et al., 2003) to obtain the MSA. We also match each sequence of one partner to the corresponding sequence of the other partner by comparing the source organisms for each protein. The assumption is that, if the proteins are homologous and interact in one organism, they should also interact in the other organisms. This is a crucial step because proteins will only coevolve within the same lineage, and without matching the correct organisms the data obtained would be meaningless. In fact, the data that can be obtained from a random match can provide a baseline with which to compare the coevolution indicators, as we explain below.

For the first test case (see Results and Discussion), the next step was to use the MSA to predict a contact map by estimating how likely it is that residues interact across a protein-protein interface. This estimate is calculated by summing all contact propensity scores (the volume-normalized scores from (Glaser et al., 2001)) for all pairs of columns in the MSA, across the two different partners. These MSA columns represent a snapshot of the evolutionary relations for a given pair of residues in all different protein sequences, and give us an indication of the overall contact propensity between those two residues across the set of proteins being considered. From this score we then subtract the total score obtained using those same two columns but randomizing the pairing of the amino acid residues. This final score is an indication of how much the contact propensity was maintained during evolution despite the occurrence of mutations that change the amino acid sequences. The higher this value, the higher the indication that compensatory mutations arose that mitigated the deleterious effects of mutations interfering with the stability of the interaction. This is a simple and intuitive measure that, though likely inferior to more refined statistical measures typically used for these purposes, still provides a suitable test to the capacity of our constrained docking algorithm to deal with noisy data. Finally, we average, for each residue in each protein, the best ten scores obtained in the pairings with all residues of the other protein. The reasoning is that residues that are at the interface must be in proximity to several residues of the other protein, and thus this average can mitigate the effect of high scores in incorrect pairs due to chance. This average also eliminates the information about which specific pairs are in contact, but this is not a problem for our system, as opposed to other constraint docking approaches that do not use CP (e.g. (Dominguez et al., 2003)), because our constraint does not require us to identify which pairs are in contact. In exchange, this averaging not only reduces the noise but also reduces by two orders of magnitude the absolute number of candidates to consider, since each protein has on the order of a hundred amino acid residues, and in this way we drop from the total number of pairs to the number of residues. This is a useful step that can only be taken because of the features of the constraint we use in docking. Applying this procedure to the first test case, we kept the five highest scoring residues after averaging, of which two were actually in the interface region. This gave a total of 85 atoms, and our constraint was that at least 20 of these 85 atoms from one partner was less than 5Å away from any atom of the other partner.

For the second test case we used CAPS (Fares and McNally, 2006) to perform a more standard coevolution analysis, based on the Pearson coefficient, and used the groups identified as coevolving to pinpoint the interaction sites. In this case the constraint is much stronger, as in each group we required at least one contact between those specific residues. This group had only two residues in one partner and one residue in the other, thus greatly reducing the number of allowed configurations. The disadvantage of this approach is that, due to the significant number of false positives in the CAPS estimates, a different docking search must be run for each group. However, given the tightness of the constraints, each docking run takes only a small fraction of the time needed for an unconstrained run, and thus all groups can be easily screened.

## 3. RESULTS AND DISCUSSION

From targets used in the Critical Assessment of Predicted Interactions experiment (CAPRI) ((Janin et al., 2003), (Janin and Wodak, 2007)) so far, we selected a complex from which coevolutionary data could, presumably, be found. This requirement excluded cases such as antibody-antigen interactions, for which no coevolution is expected, and complexes determined for the interaction of only a domain or fragment of a larger protein, since this could create additional difficulties for the MSA analysis. Of the remaining candidates, we opted for the complex formed by the bacterial N5-glutamine Methyltransferase (HemK) and Peptide Chain Release Factor 1 (RF1) (Graille et al., 2005), corresponding to target 20 in the sixth round of CAPRI. Figure 1 (panels A, B and C) summarizes the results obtained for this complex, with the docking simulations starting from randomly oriented partners.

**Figure 1:**
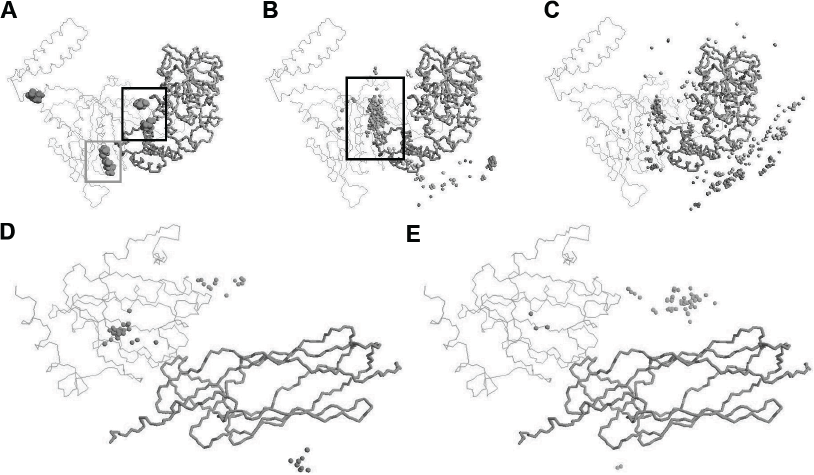
Results for the complexes RF1-HemK and DsbD-CcmG. Panel A shows the residues of RF1 identified as candidates for the interface with HemK. Two residues, in the black rectangle, were correctly identified. The remaining three are false positives, even though the perspective makes it seem that the two residues in the grey rectangle are close to the interface. Panel B shows the pattern of models obtained with the constraint, and the panel C shows the distribution of models obtained without this constraint. Panel D shows the pattern of models obtained with the constraint given by the CAPS analysis. Panel E shows the distribution of models obtained without this constraint. The lighter shaded structure shows the correct position of the partner that is represented by the spheres. The 50 highest-scoring models according to BiGGER’s contact propensity score are represented in each panel.

Panel A shows five residues of RF1 selected as candidates for the interface. The two residues inside the black rectangle are actually at the interface, and the other three are incorrect. The two residues inside the gray rectangle appear to be close to the interface due to the perspective, but are actually more than 12Å away from HemK. Figure 1B shows the correct RF1-HemK complex and, around the HemK protein, 400 spheres indicating the geometric center (average of all atomic coordinates) of the RF1 protein in each of the 400 models that scored highest in surface contact and which respected the constraint on the contacts between the selected RF1 residues and any atom in HemK. The spheres inside the black rectangle indicate the large cluster of models close to the correct position. It is important to note that this constraint only specified a set of residues on RF1. For HemK, any atom could count for a contact. However, this was enough not only to place HemK in the correct position relative to RF1 but also, in conjunction with the surface contact criteria, to place RF1 in the correct position relative to HemK in a large number of models. The panel C shows the best 400 models without this contact constraint obtained from the five candidate residues. It is easy to see that, in this case, the positions of RF1 more widely distributed around HemK.

Comparing these two sets of 400 models with the largest surface contact, the set without the constraint contains 2 models similar to the correct complex (¡5Å of RMSD) and a median RMSD of 26Å when compared to the correct complex. In contrast, the set obtained by imposing the constraint has 5 models similar to the actual complex and a median RMSD of 17.5Å. This is a significant improvement because the greatest problem in protein-protein docking is to identify the correct models within the set of models that have the best surface contact (Halperin et al., 2002). Thus, anything that helps to enrich this set with good approximations of the target complex makes this task easier.

Our other target complex was the N-termimal region of the Disulfide interchange protein (DsbD) and Cytochrome c biogenesis protein (CcmG) (Stirnimann et al., 2005), from the protein-protein docking benchmark (Hwang et al., 2008). For this complex we used CAPS, a software package designed to perform coevolution analysis on protein sequences with improved sensitivity over previous methods for detecting intra-molecular coevolution, and also capable of detecting inter-molecular coevolution. CAPS uses Blosum-corrected amino acid distances to identify co-variation between residues across different proteins and measures their correlation using the Pearson coefficient ((Fares and McNally, 2006), (Fares and Travers, 2006)). CAPS can thus provide estimates on groups of residues that coevolve, and these groups can be used to greatly constrain the search space for permitted configurations. It is important to note that these estimates are noisy and, in general, a large part of the groups identified by CAPS are false positives. However, the constraints provided by forcing contacts between small groups of residues reduce the computation time for each docking run from several hours to around ten minutes. This makes it practical to test all the ten to twenty groups that CAPS typically proposes as coevolving in even less time that it would take to run the docking search without constraints. As Figure 1 (panels D and E) illustrates, the results are quite significant. On the panel D we show the best 50 models, as scored by BiGGER according to residue contact propensities, from the constrained docking run. The panel E shows also the best 50 models, but from an unconstrained run. Whereas in the unconstrained docking there are only 3 models in the correct position, as shown by the three spheres inside the lighter structure representing the position of the partner in the target complex, in the constrained docking more than half of the highest-scoring models are close to the correct position, greatly simplifying the task of adequately modeling the structure of this complex. This means that it is feasible to test, individually, all potential residue contacts given by the coevolution measurements. Furthermore, since the docking runs are independent, the whole process is trivial to run in parallel, thus, in practice, taking less time than a single unconstrained docking run.

## 4. CONCLUSIONS

The work presented here leaves many questions still unanswered. There are still many parameters to optimize, such as the number of candidates to consider, the thresholds for the contact distances and the number of contacts, as well as implementation details, such the integration of better scoring functions for the inference of coevolution, and also the need to apply this approach to other test cases and gather more data on its advantages and shortcomings. However, these tasks are ongoing, part of current and intended future work. For the work reported here, the purpose was to test whether CP techniques that enforce strict and well defined constraints (such as our cardinality constraint on the set of contacts) can be profitably applied to help processing diffuse and noisy statistical data. Our test cases suggest that this is true and thus, we hope, can motivate more applications of CP in those areas of Bioinformatics traditionally dominated by “softer”, more statistically oriented, approaches.

The software referred to and developed as part of this work is available, included in the Open Chemera Library (Krippahl, 2012).

## ACKNOWLEDGMENTS

This work was funded by FEDER, COMPETE and FCT, under project PTDC/EIA-CCO/115999/2009. We also thank Marco Correia and Pedro Barahona for their suggestions and discussions during this work.

## REFERENCES

Altschul, S. F., Madden, T. L., Schäffer, A. A., Zhang, J., Zhang, Z., Miller, W., Lipman, D. J. (1997). Gapped BLAST and PSI-BLAST: a new generation of protein database search programs. Nucleic Acids Research, 25(17), 3389–402.

Chenna, R., Sugawara, H., Koike, T., Lopez, R., Gibson, T. J., Higgins, D. G., Thompson, J. D. (2003). Multiple sequence alignment with the Clustal series of programs. Nucleic Acids Research, 31(13), 3497–500.

Codoñer, F.M., Fares, M.A. (2008). Why should we care about molecular coevolution? Evolutionary Bioinformatics, 4, 29–38.

Dominguez C., Boelens R., Bonvin A.M. (2003). HADDOCK: a protein-protein docking approach based on biochemical or biophysical information. Journal of the American Chemical Society, 125(7), 1731–1737.

Elias, I. (2006). Settling the intractability of multiple alignment. Journal of Computational Biology, 13(7), 1323–39.

Fares M.A., McNally D. (2006). CAPS: coevolution analysis using protein sequences. Bioinformatics, 22(22), 2821–2822.

Fares M.A., Travers S.A. (2006). A novel method for detecting intramolecular coevolution: adding a further dimension to selective constraints analyses. Genetics, 173, 9–23.

Glaser F., Steinberg D.M., Vakser I.A., Ben-Tal N. (2001). Residue frequencies and pairing preferences at protein-protein interfaces. Proteins, 43(2), 89–102.

Graille M., Heurgué-Hamard V., Champ S., Mora L., Scrima N., Ulryck N., van Tilbeurgh H., Buckingham R.H. (2005). Molecular basis for bacterial class I release factor methylation by PrmC. Molecular Cell, 20(6), 917–927.

Halperin I., Ma B., Wolfson H., Nussinov R. (2002). Principles of docking: An overview of search algorithms and a guide to scoring functions. Proteins, 47(4), 409–443.

Hwang H., Pierce B., Mintseris J., Janin J., Weng Z. (2008). Protein-protein docking benchmark version 3.0. Proteins, 73(3), 705–709.

Halperin, I., Wolfson, H., Nussinov, R. (2006). Correlated Mutations: Advances and Limitations. A Study on Fusion Proteins and on the Cohesin-Dockerin Families. Proteins: Structure, Function, and Bioinformatics, 63(4), 832–845.

Horner, D. S., Pirovano, W., Pesole, G. (2007). Correlated substitution analysis and the prediction of amino acid structural contacts. Briefings in Bioinformatics, 9(1), 46–56.

Janin J., Henrick K., Moult J., Eyck L.T., Sternberg M.J., Vajda S., Vakser I., Wodak S.J. (2003). CAPRI: a Critical Assessment of PRedicted Interactions. Proteins, 52, 2–9.

Janin J., Wodak S. (2007). The third CAPRI assessment meeting Toronto, Canada, April 20-21, 2007. Structure, 15(7), 755–759.

Just W. (2001): Computational complexity of multiple sequence alignment with SP-score. Journal Computational Biology, 8(6), 615–623.

Krippahl, L. (2012). Open Chemera Library. Available at https://github.com/lkrippahl/Open-Chemera

Krippahl L., Barahona P. (2005). Applying Constraint Programming to Rigid Body Protein Docking. Principles and Practice of Constraint Programming-CP 2005. 373–387.

Krippahl L., Moura J.J., Palma P.N. (2003). Modeling protein complexes with BiGGER. Proteins, 52, 19–23.

Little D.Y., Chen L. (2009). Identification of Coevolving Residues and Coevolution Potentials Emphasizing Structure, Bond Formation and Catalytic Coordination in Protein Evolution. PLoS ONE, 4(3), e4762.

Lovell, S. C., Robertson, D. L. (2010). An integrated view of molecular coevolution in protein-protein interactions. Molecular Biology and Evolution, 27(11), 2567–2575.

Madaoui H., Guerois R. (2008). Coevolution at protein complex interfaces can be detected by the complementarity trace with important impact for predictive docking. Proceedings National Academy of Science, 105(22), 7708–7713.

Monaco S., Gioia M., Rodriguez J., Fasciglione G.F., Pierro D.D., Lupidi G., Krippahl L., Marini S., Coletta M. (2007). Modulation of the proteolytic activity of matrix metalloproteinase-2 (gelatinase A) on Fibrinogen. Biochemical Journal, 402(3), 503–513.

Myers G., Selznick S., Zhang Z., Miller W. (1996). Progressive Multiple Alignment with Constraints. Journal Computational Biology, 3, 563–572.

Papadopoulos J.S., Agarwala R. (2007). COBALT: constraint-based alignment tool for multiple protein sequences. Bioinformatics, 23(9), 1073–1079.

Pazos, F., Helmer-Citterich, M., Ausiello, G., Valencia, A. (1997). Correlated mutations contain information about protein-protein interaction. Journal of Molecular Biology, 271(4), 511–23.

Pazos, F.,Valencia, A., (2008). Protein co-evolution, co-adaptation and interactions. The EMBO Journal, 27(20), 2648–55.

Palma, P. N., Krippahl, L., Wampler, J. E., Moura, J. J. (2000). BiGGER: a new (soft) docking algorithm for predicting protein interactions. Proteins, 39(4), 372–84.

Palma P.N., Lagoutte B., Krippahl L., Moura J.J., Guerlesquin F. (2005). Synechocystis ferredoxin/ferredoxin-NADP(+)-reductase/NADP + complex: Structural model obtained by NMR-restrained docking. FEBS Letters, 579(21), 4585–4590.

Sammeth M., Morgenstern B., Stoye J. (2003). Divide-and-conquer multiple alignment with segment-based constraints. Bioinformatics, 19 Suppl 2, ii189–ii195.

Stirnimann C.U., Rozhkova A., Grauschopf U., Grütter M.G., Glockshuber R., Capitani G. (2005). Structural basis and kinetics of DsbD-dependent cytochrome c maturation. Structure, 13(7), 985–993.

Wang L., Jiang T. (1994). On the complexity of multiple sequence alignment. Journal Computational Biology, 1(4), 337–348.

Will S., Busch A., Backofen R. (2008). Efficient Sequence Alignment with Side-Constraints by Cluster Tree Elimination. Constraints Journal, 13, 110–129.

Yap R. (2001). Parametric Sequence Alignment with Constraints. Constraints Journal, 6, 2001.

Yeang C.H., Haussler D. (2007). Detecting coevolution in and among protein domains. PLoS Computational Biology, 3(11), e211.

